# Stem cell growth directs region-specific cell fate decisions during intestinal nutrient adaptation

**DOI:** 10.1101/2023.04.24.537654

**Authors:** Jaakko Mattila, Arto Viitanen, Gaia Fabris, Jerome Korzelius, Ville Hietakangas

**Affiliations:** Faculty of Biological and Environmental Sciences, University of Helsinki, Helsinki 00790, Finland; Institute of Biotechnology, University of Helsinki, Helsinki 00790, Finland; School of Biosciences, University of Kent, Canterbury, CT2 7NJ, UK

## Abstract

The adult intestine is a regionalized organ, whose size and cellular composition is adjusted in response to nutrient status. This involves dynamic regulation of intestinal stem cell (ISC) proliferation and differentiation. How nutrient signaling controls cell fate decisions to drive regional changes in cell type composition remains unclear. Here we show that nutrient adaptation involves region-specific control of intestinal cell size, number and differentiation. We uncovered that activation of mTOR complex 1 increases ISC size in a region-specific manner. This promotes Delta expression to direct cell fate towards the absorptive enteroblast lineage, while inhibiting secretory enteroendocrine cell differentiation. The observed coupling between nutrient sensing and cell fate enabled mitigation of aging-induced ISC misdifferentiation through intermittent fasting. In conclusion, ISC size acts as an early fate determinant allowing regional control of intestinal cell differentiation in response to nutrition with relevance to maintenance of tissue integrity during aging.

**Highlights:**

- mTORC1 signaling regulates ISC size in a region-specific manner
- mTORC1 signaling is activated in the S and G2 phase of the ISC cell cycle
- ISC size directs differentiation towards absorptive vs. secretory lineage
- Intermittent fasting mitigates aging induced deregulation of ISC differentiation

**GRAPHICAL ABSTRACT:** 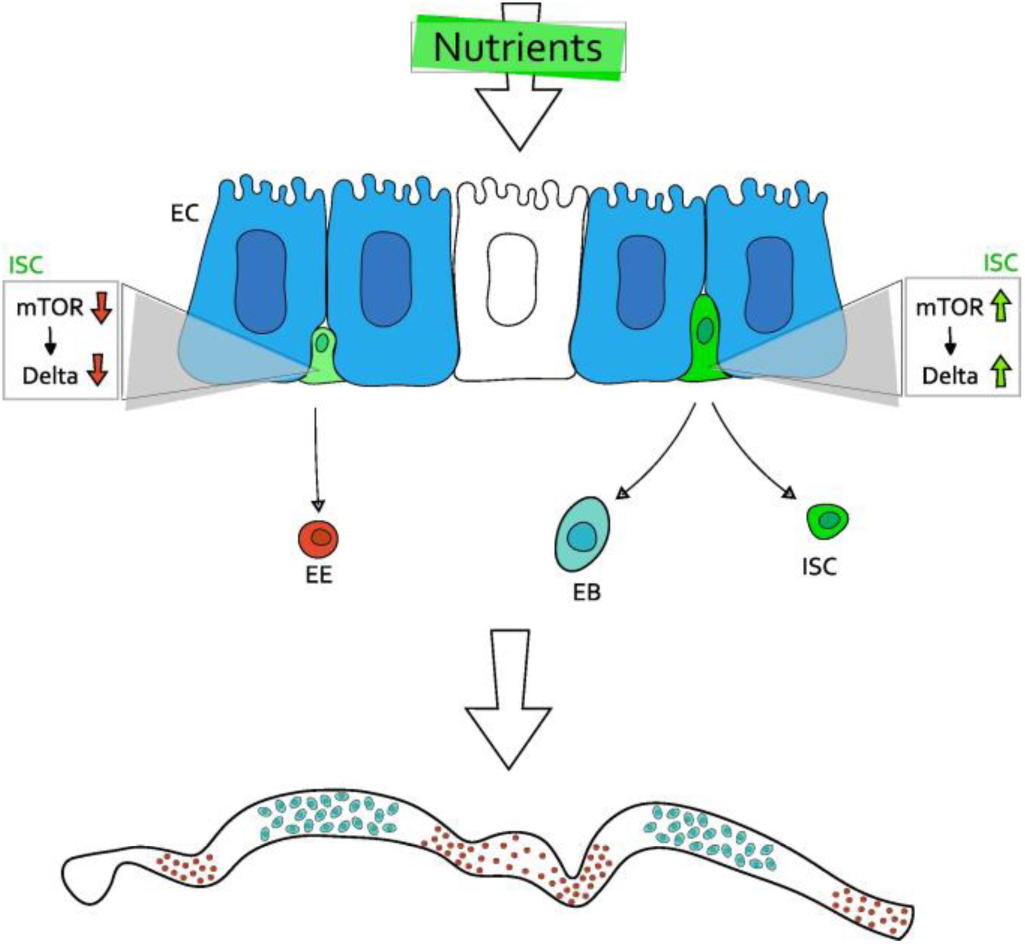

## INTRODUCTION

Nutrient intake modulates the physiology of adult animals through alteration of metabolism of tissues, but also through changes in cellular composition of organs through the activity of somatic stem cells. For example, the net volume and morphology of the small intestine is strongly regulated by nutrition to match the organ’s absorptive, metabolic and signaling functions with the physiological needs of the animal^1–3^. This includes coordinated control of proliferation and differentiation of stem cells and size of differentiated cells^2–4^. How the balance between these parameters is dynamically controlled by nutrient signaling in the spatial context of organs, for example in different intestinal regions, remains poorly understood.

The *Drosophila* midgut, the counterpart of mammalian small intestine, contains four distinct cell types: large absorptive enterocytes (ECs) and their precursors enteroblasts (EBs), as well as the small secretory enteroendocrine (EE) cells, all differentiating from the mitotic intestinal stem cells (ISCs)^5^. The EB/EC and EE differentiation is determined by the strength of Notch signaling. ISCs with high Delta expression direct their daughter cells towards EB/EC fate while ISCs with low Delta promote EE differentiation^6,7^. Feeding of experimental diet with high cholesterol reduces Delta-Notch signaling, promoting ISC differentiation towards the EE cell fate^8^. However, whether the adaptive growth of the intestine upon transition from fasted to fed state ^2^ involves differential control of EB/EC and EE fates remains to be addressed. The ISC differentiation towards the EB/EC lineage involves a prominent increase in cell size, which is mediated by Notch-dependent activation of mTORC1 in enteroblasts, driving EB differentiation towards the EC fate^9^.

Midgut size is highly adaptive to nutrition: When calorie-restricted, ISC proliferation is low and the midgut volume decreases through EC loss and size reduction^2,4^. Upon feeding, the proliferation and differentiation of ISCs increased and the ECs gain in size, leading to increase in midgut volume^2,4,10,11^. This nutrient-induced ISC proliferation depends on insulin-induced PI3K/AKT signaling^2,11^, while the EC size increase upon feeding requires mTORC1 activity^4^. Both mammalian and *Drosophila* intestines are highly regionalized organs^12–14^. The *Drosophila* midgut is divided into six (R0-R5) anatomically recognizable regions with characteristic gene expression patterns and cellular content^12,14–16^. The regulation of ISCs is spatially defined, as evidenced by region-specific enrichment of ISC daughter cells upon DSS-induced tissue damage response^16^. However, our current understanding on the regional specificities of nutrient regulation of intestinal cells remains very limited.

The mTOR complex 1 (mTORC1) is a key regulator of nutrient-induced growth in differentiated cells and tissues^17^. However, the role of mTORC1 signaling in somatic stem cells is more complex and context-dependent. Several lines of evidence imply that sustained mTORC1 signaling impairs stem cell function. In epithelial stem cells constitutive mTORC1 activation by Wnt signaling leads to increased growth of hair follicles and disappearance of the epidermal stem cell compartment^18^. In *Drosophila* ISCs high mTORC1 activity upon TSC1 and TSC2 loss-of-function leads to massively increased cell size, but inhibits proliferation and differentiation^19^. Moreover, aging increases the size of hematopoietic stem cells (HSCs) impairing their function, which can be rescued by inhibition of mTORC1 signaling^20^. In contrast to sustained mTORC1 activity, short-term stimulation of mTORC1 signaling does not compromise stem cell function and is involved in their physiological regulation. In quiescent *Drosophila* ISCs, mTORC1 activity is inhibited by high expression of TSC2^9^. Upon tissue damage-induced regeneration mTORC1 signaling is transiently activated in the ISCs, which is necessary for ISC proliferation and consequent intestinal regeneration^21^. Repeated cycles of ISC mTORC1 activation, however, lead to stem cell loss^21^. In addition, mTORC1 is necessary to reactivate mouse muscle stem cells from quiescence after injury^22^. In the intestine of nutrient-restricted mice, mTORC1 signaling in neighboring Paneth cells is inhibited, enhancing ISC function^23^. How the nutrient-dependent mTORC1 signaling influences the proliferation and lineage differentiation decisions in of adult stem cells remains poorly understood.

Here we use an organ-wide approach to quantitatively analyze the regulation of cell size, number and identity upon nutrient-induced tissue adaptation of the *Drosophila* midgut. Our data reveals a striking regional heterogeneity in the regulation of EC growth, as well as ISC division and differentiation by nutrient-induced cues. Following ISC activation upon acute transition from fasting to feeding increases the number of EBs and EE cells, but with contrasting regional distributions. We also observed an mTORC1-mediated increase in ISC size, which was particularly strong in the regions with nutrient-induced EB accumulation. Consistent with the regional distributions of EB and EE cells, this ISC growth primed them to divide asymmetrically producing more EBs, while inhibiting the differentiation towards EE cell lineage. ISC mTORC1 activity promoted high expression of Delta, which is known to facilitate EB differentiation in recently divided ISC-EB pairs. In aged animals the ISCs respond poorly to feeding and display abnormally elevated ISC and EE fate, compared to young animals. This aging-induced deregulation of ISC differentiation can be, however, mitigated by intermittent fasting. Collectively our data demonstrates that stem cell size can act as an early fate determinant to mediate region-specific differentiation patterns upon intestinal nutrient adaptation with relevance to prevention of aging-associated tissue decline.

## RESULTS

### Organ-wide analysis reveals regional heterogeneity of midgut adaptive growth

Transition from fasted to fed state promotes ISC proliferation and enterocyte growth, enabling dynamic adjustment of adult midgut size^2,4,24^. Current knowledge on midgut growth regulation relies on local quantifications of distinct midgut regions, lacking the global organ-wide insight to address a possible role for region-specific regulation. Consistent with previous findings, feeding of flies on complete holidic diet after fasting on a highly restricted diet (2% sucrose) stimulated midgut growth and increased total cell numbers on both sexes (Figure 1A, B; Figure S1A-F). To achieve organ-wide insight into this adaptive growth regulation, we utilized the recently-developed image analysis method, LAM, that allows region-specific analysis of cellular parameters of the intestine^16^ (Figure 1C). Refeeding led to >1.5-fold increase in midgut width (Figure 1D), while largely retaining the morphology of the regional boundaries (Figure 1E), allowing us to align the intestinal regions of midguts with distinct size.

**Figure 1.**
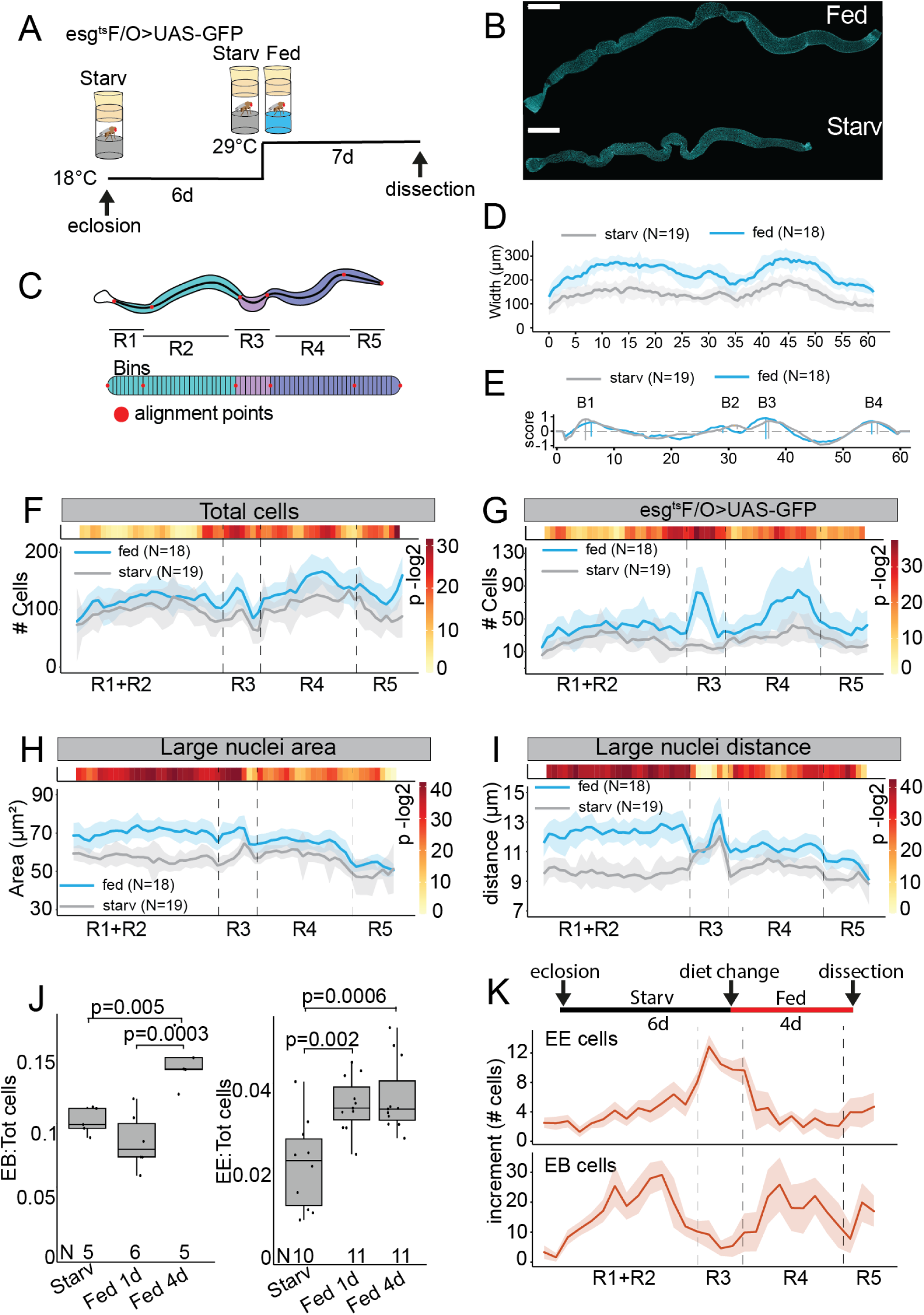
Midgut nutrient adaptation is regionally defined. **A)** Experimental design used to obtain data in panels B-I. Age matched, mated esg^ts^F/O>UAS-GFP females, (eclosed to starvation diet, 2% sucrose), were kept at +18°C for six days, and then shifted to the permissive temperature (+29°C) for additional 7 days in either starvation or holidic diet. **B)** Representative images of DAPI (cyan) stained midguts of females kept in either holidic diet or in starvation diet. Scale bar 200 μm. **C)** Principle of LAM. LAM transforms image-derived cellular data from three-dimensional midguts into a linearized representation, binning it into segments along the A/P axis. As a result, LAM allows region to region comparison of midguts of starved and fed animals. **D**) Feeding results in regionally uniform midgut growth. Width profile of starved and fed midguts along the midgut A/P axis. **E)** Region border analysis show little variation in the positions of region borders between starved and fed female midguts. The marked borders from left to right are B1, B2, B3, and B4. **F-I)** Feeding induces regionally distinct pattern of cell proliferation and cell growth. Total cell counts **(F)**, GFP positive cell counts **(G)**, large nuclei area **(H)** and nearest distance between large nuclei **(I)** along the A/P axis of starved and fed female midguts of genotype esg^ts^F/O>UAS-GFP. Dashed lines indicate the main region borders. Light blue/grey shading is the standard deviation. **J-K)** Feeding induces temporally and spatially defined increase in EE and EB cells. **J)** Relative number of EB and EE cells in starved, one day fed and four days fed midguts. **K)** Increase of EE and EB cell numbers along the midgut A/P axis four days after commencement of feeding. P values in **F-I** were obtained by Wilcoxon rank-sum test using continuity correction. In the bin-by-bin testing, false discovery rate correction due to multiple testing is applied. P values in **J** were obtained by two-way ANOVA followed by Tukey’s test. See also Figure S1.

Despite the relatively uniform increase in midgut width, quantitative analysis of specific cellular parameters uncovered striking regional variation in the mode of nutrient regulation. An increase in total cell numbers was observed in specific areas of the central and posterior regions, while R1 and R2AB regions displayed limited increase in cell numbers (Figure 1F). Consistently, an increase in stem cell activity, as measured by the GFP-marked cells in esg^ts^ Flp-Out (esg^ts^F/O) clones^25^, was highest in the regions with increased cell numbers, including anterior R3 (Copper Cell Region) and in R4bc, whereas the anterior regions were less affected (Figure 1G). In contrast, EC size was most increased in the R1 and R2 regions (Figure 1H,I). As a surrogate to cell size, we measured the maximum nucleus cross section area (hereafter referred as nuclear area) from midguts of starved and fed flies. The nuclear area correlates well with cell size, and is amenable to automated quantification from 3D-segmented images of DAPI stained nuclei (Figure S1G,H). Elimination of stem cells by Reaper overexpression revealed that the adaptive intestinal growth can occur near-normally, consistent with earlier findings (Figure S1I-K)^4^.

Next, we wanted to address in detail the regional increase in cell numbers. Indeed, feeding induced specific changes in the numbers of ISC daughter cells, i.e. EBs and EE cells. While the relative number of both cell types was increased upon feeding, the response occurred through distinct kinetics. Indeed, EE cell numbers increased after 1 day of feeding, while EBs numbers increase 4 days after feeding (Figure 1J). Strikingly, we observed opposite regional pattern in the distribution of EB and EE number increase. While EE cell numbers increased mainly in R3 and the region borders flanking R3, EB numbers were mostly elevated in R2 and R4 regions (Figure 1K). Collectively, our organ-wide analysis uncovered regional differences in cell type composition and ISC proliferation as response to nutrient availability in the midgut.

### Region-specific mTORC1 activation controls ISC size upon feeding

Organ-wide analysis of the size profiles of all intestinal cells showed that feeding-induced growth is pervasive, as it can be observed at the level of the whole population of polyploid ECs (Figure 2A). Surprisingly, the global analysis of cellular size distributions also revealed a size increase of the small diploid cells, which include the ISCs (Figure 2A). Therefore, we specifically analyzed the feeding-induced size regulation of Delta^+^ ISCs (Figure 2B, C). Indeed, ISC size increased significantly upon feeding, being most prominent already at the 1-day time point (Figure 2C, D). Thus, ISC enlargement preceded the increase in EB cell numbers (Figure 1J). Next, we analyzed whether the ISC size is regionally regulated upon changing nutrient intake. Nutrient-induced ISC growth was highly regionalized in starved and fed animals: ISCs at R2, R4 and R5 displayed feeding-induced size increase, while ISC size in R1 as well as the R2-R3 and R3-R4 borders regions were unresponsive to nutrient availability (Figure 2E & F). Thus, the feeding-induced control of ISC size is region specific. Interestingly, the spatial distribution of ISC growth corresponded to the distribution of EB accumulation upon feeding (Figure 1K).

**Figure 2.**
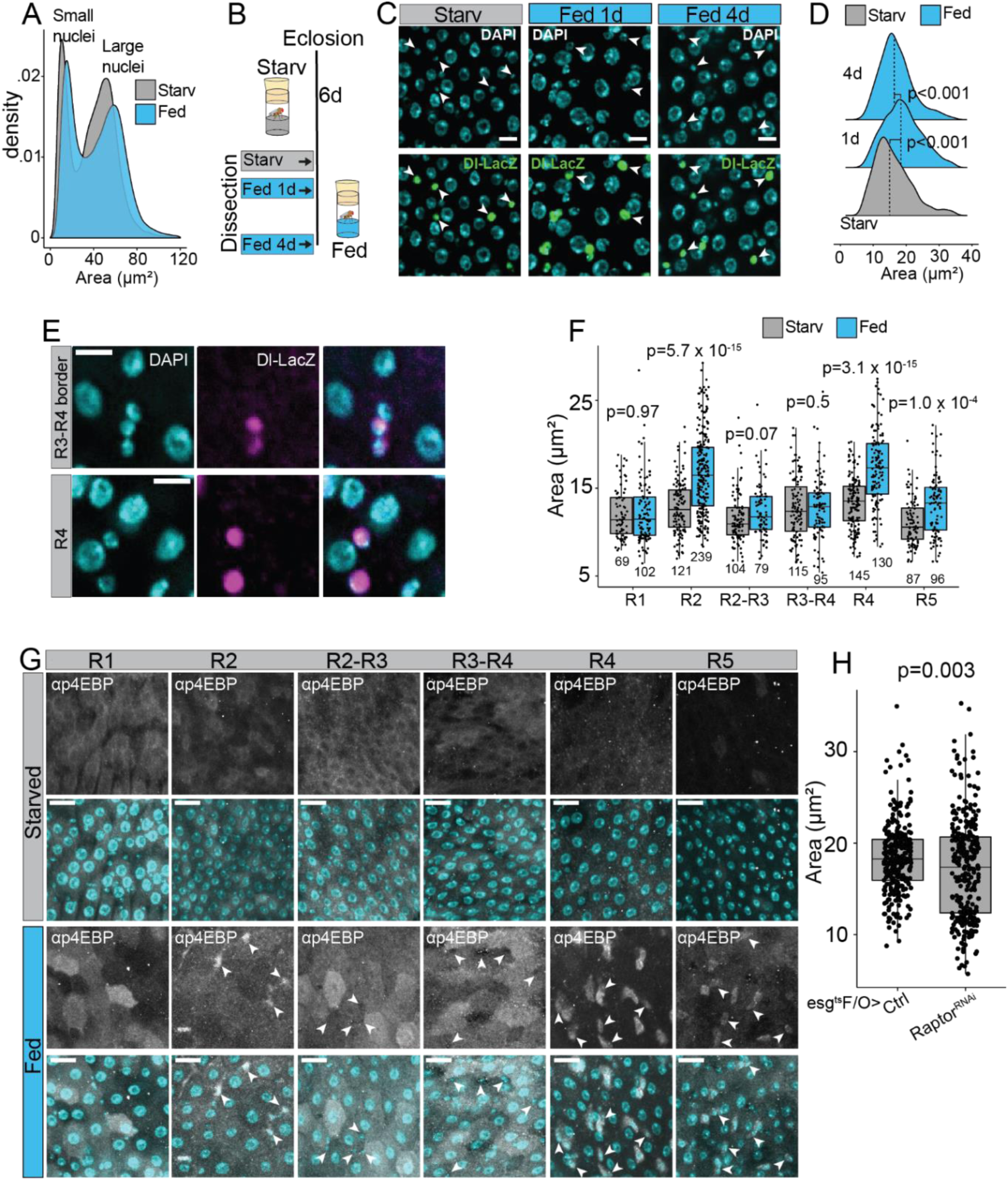
ISC size is regulated regionally by feeding-induced mTORC1 signaling. **A)** Nucleus area distribution in starved versus fed female midguts. The size of small and large nuclei is increased in midguts of fed flies. Pooled data from N^starved^=12 and N^fed^=12 full midgut images, and N^starved^=64477 and N^fed^=87505 cells. **B)** Experimental design used to obtain data in panels C & D. Age matched, mated females harboring the Delta-LacZ marker were aged for six days in starvation (2% sucrose), and then shifted to the holidic diet for an additional 1 or 4 days. **C-D)** The size of ISC nuclei is temporally regulated in midguts of fed flies. **C)** Representative images from Delta-LacZ female midguts stained with α-β-Galactosidase (green) and DAPI (cyan). Arrowheads point to ISCs positive for Delta-LacZ marker. Scale bar 10μm. **D)** Quantification of Delta-LacZ positive ISC nucleus area from the experiment depicted in B & C. Quantifications were made from the R4b region. **E-F)** ISC size is regulated regionally by feeding. **E)** Representative images of ISCs (marked by α-β-Galactosidase staining of Delta-LacZ) in the R4 and R3-R4 border regions. Scale bar 10μm. **F)** Quantification of nucleus area of Delta-LacZ positive ISCs from R1, R2, R2-R3 border, R3-R4 border, R4 and R5 regions from starved and fed female midguts. Pooled data from N^starved^=3, N^fed^=3 midguts. N^cells^ are indicated in the figure panel. **G-H)** mTORC1 activation by feeding is region- and cell-type-specific. **G)** Representative images of midgut regions from female flies kept in either starvation or holidic diet and stained by αp4EBP (grey) and DAPI (cyan). Experimental design as in Figure 1A. Scale bar 20μm **H)** Quantification of Delta-LacZ positive ISC nucleus area from esg^ts^F/O>UAS-Raptor^RNAi^, Delta-LacZ. p values in **D** were obtained by two-way ANOVA followed by Tukey’s test. p values in **F** were obtained by Wilcoxon rank-sum test with multiple testing correction (FDR<0.05). p value in **H** is calculated by the two-sample t-test.

To understand how the region-specific ISC growth is regulated, we investigated the role of the mTORC1 signaling pathway, a well-known regulator of cell size^17^. Midguts were analyzed for the mTORC1 target 4EBP phosphorylation (p4EBP), in the regions R1, R2, R4, R5 as well as the region borders flanking R3. Consistent with its role as cellular nutrient sensor, mTORC1 activity showed increased activity in response to feeding in all intestinal regions (Figure 2G). However, the cellular distribution of mTORC1 activity displayed striking regional heterogeneity. In the R1 region, as well as in the borders flanking R3, p4EBP signal is mainly detected in polyploid ECs, while in R4 and R5 p4EBP signal is high in the small nuclei cell population (Figure 2G). In R2, p4EBP signal is mixed, showing signal in both large and small cells. To explore the functional importance of mTORC1 signaling in the ISCs, we depleted mTORC1 activity by RNAi knockdown of Raptor using the esg-Gal4 driver and measured nuclear area from Delta-LacZ positive ISCs. Indeed, ISCs from fed animals with reduced mTORC1 signaling were significantly smaller in size as compared to the controls (Figure 2H). In conclusion, ISCs display a region-specific increase in size as an immediate response to feeding, which is controlled by mTORC1 activation.

### Cell cycle-specific regulation of ISC mTORC1 signaling

To further analyze mTORC1 activity in the diploid cell population, we explored the p4EBP signal pattern specifically in the Delta^+^ ISC population. The mTORC1 activity was heterogeneous among ISCs. We identified both single ISCs and ISC-ISC doublets, either positive or negative for the mTORC1 marker p4EBP (Figure 3A-C). This heterogeneity led us to test whether mTORC1 activity depends on the cell cycle phase. We used Delta-Gal4 to express the fluorescent cell cycle specific FUCCI reporter (fly-FUCCI)^26^ in the ISCs, which allowed us to analyze 4EBP phosphorylation during G1, S and G2/M phases separately. This analysis revealed that mTORC1 activity is low in G1 and gets gradually elevated in S and G2/M phases (Figure 3D, E), consistent with the observed heterogeneity of mTORC1 activity in the total ISC population. We also used the fly-FUCCI system to analyze feeding-induced ISC growth. Our data shows that ISCs of the fed animals were significantly larger compared to the starved animals at all cell cycle phases and the difference between the size distributions gradually increased in S and G2/M phases, consistent with the mTORC1 activity (Figure 3F & 3G). Thus, the mTORC1 dependent ISC growth is gradually activated while the cell cycle progresses towards mitosis.

**Figure 3.**
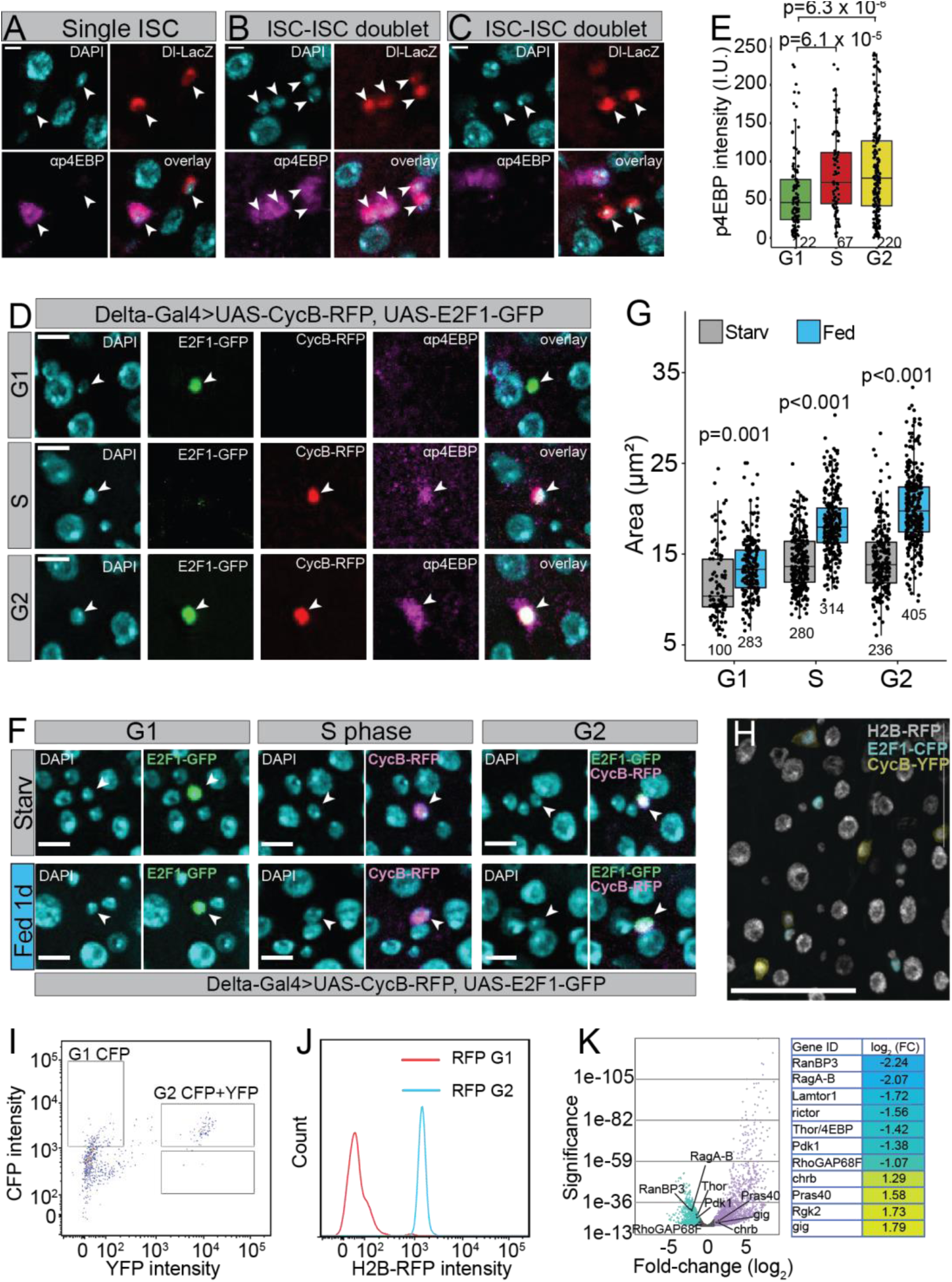
ISC mTORC1 activity is regulated in a cell cycle dependent manner. **A-C)** mTORC1 activity is heterogeneous between ISCs. Representative images of single **(A)** and doublet **(B & C)** ISCs stained by αp4EBP (magenta), αβ-Galactosidase (red), and DAPI (cyan). Images are from the R4b region of Delta-LacZ ISC marker bearing female flies kept in holidic diet for 7 days. Scale bar 5μm. **D-E)** mTORC1 activity is elevated in the S and G2/M phases of the cell cycle. **D)** Representative images from midguts of Delta-Gal4>UAS-CycB-RFP, UAS-E2F1-GFP (fly-FUCCI) stained by αp4EBP (magenta) and DAPI (cyan). Scale bar 10μm. **E)** Background subtracted ISC nucleus p4EBP intensity from G1, S and G2 phase from experiment depicted in **D**. Pooled data from N=4 midguts from the R4b region. N^cells^ are indicated in the figure panel. **F-G)** ISCs of the fed animals are larger compared to the starved animals at all cell cycle phases. **F)** Representative images of G1, S and G2/M phase ISCs from midguts of female flies of genotype Delta-Gal4>UAS-CycB-RFP, UAS-E2F1-GFP (fly-FUCCI) kept in either starvation or holidic diet. DAPI (cyan), E2F1-GFP (green) and CycB-RFP (magenta). Scale bar 10 μm **G)** Quantification of the experiment depicted in **F**. Pooled data from N^starved^=3, N^fed^=4 midguts. N^cells^ are indicated in the figure panel. **H-K)** Gene expression profiling of G1 vs. G2/M ISCs show differential expression of mTORC1 regulators. **H)** Representative image of the UAS-driven CFP::E2F1/YFP::NLS-CycB transgenic Fly-FUCCI line (cyan and yellow, respectively) combined with His2AV-mRFP1 (gray). Fly-FUCCI is driven specifically in ISCs using the ISC-specific esg-Gal4, Su(H)GBE-Gal80 driver line. **I)** Gating strategy to sort G1 (CFP only) and G2 (CFP+YFP) cells. **J)** An extra gating check was made by counting the H2B::RFP intensity for the two different populations. Two clear peaks, G1 (red) and G2 (blue), were detected. **K)** mTOR-associated genes enriched and depleted in G1 stem cells. Positive mTORC1 regulators (ragA-B, Lamtor1, Pdk1) are depleted in G1 ISCs, whereas negative regulators (gig/TSC2, PRAS40, charybdis) are upregulated in G1 ISCs. p values in **E** and **G** were obtained by two-way ANOVA followed by Tukey’s test. See also Figure S2.

To better understand the cell cycle specific regulation of mTORC1 activity in the ISCs, we performed gene expression profiling in G1 vs. G2/M ISCs, employing FACS-sorting of FUCCI-marked ISCs. Briefly, we combined a UAS-driven CFP::E2F1/YFP::NLS-CycB fly-FUCCI with His2AV::mRFP (to mark all nuclei) and used the ISC-specific esg-Gal4, Su(H)GBE-Gal80 line to restrict expression to ISCs (Figure 3H). The CFP and YFP signals were used to separate G1 and G2/M ISCs and total RNA was isolated from both populations (Figure 3H, I). Analysis of the His2AV-mRFP fluorescence signal in G1 and G2/M populations shows a clear distinction of these populations based on nuclear His2AV levels (Figure 3J). Differential Expression (DE) analysis identified 1690 genes upregulated and 1235 genes downregulated in G1 ISCs compared to G2/M ISCs. Among the downregulated genes, we found several positive regulators of mTORC1-activity, such as ragA-B, Lamtor1, and Pdk1 (Figure 3K). Conversely, we found G1 specific upregulation of several negative mTORC1 regulators, such as charybdis, gigas (gig/TSC2) and PRAS40^27^. Taken together, our comparison of the transcriptomes of ISCs in G1 and G2/M phases of the cell cycle, suggest that the ISCs in G1 keep mTORC1 in an inhibited state, which is released upon cell cycle progression to G2/M.

### ISC mTORC1 signaling directs cell fate

What is the role of mTORC1-mediated growth in ISCs? As the mTORC1 activity was strongest in the G2/M phase, we hypothesized that high mTORC1 activity might influence the mode of cell division. ISCs can divide either symmetrically to produce two ISCs or asymmetrically into one ISC and one EB, which undergoes rapid growth during differentiation towards EC fate^2^. Therefore, we wanted to explore the possibility that high mTORC1 activity in ISC directs cells to towards the EB fate through asymmetric division. As we were not able to directly follow the dynamic regulation of mTORC1 activity prior to symmetric vs. asymmetric division, we quantified the size of ISCs in recently divided ISC-ISC vs. ISC-EB doublets. This was based on the assumption that the growth regulation prior to division is reflected to the size of the newly divided ISCs. Indeed, the mean ISC size in ISC-ISC doublets was substantially smaller when compared to ISCs present in asymmetric ISC-EB doublets (Figure 4A). We also scored the number of Delta-LacZ positive ISCs present in esg^+^ cell doublets, and cell triplets from midguts of either starved or fed animals (Figure S2A). Consistent with the model of nutrient availability in favoring asymmetric ISC-EB division, the proportion of doublets and triplets containing two ISCs was higher in midguts of starved animals compared to the fed ones (Figure 4B).

**Figure 4.**
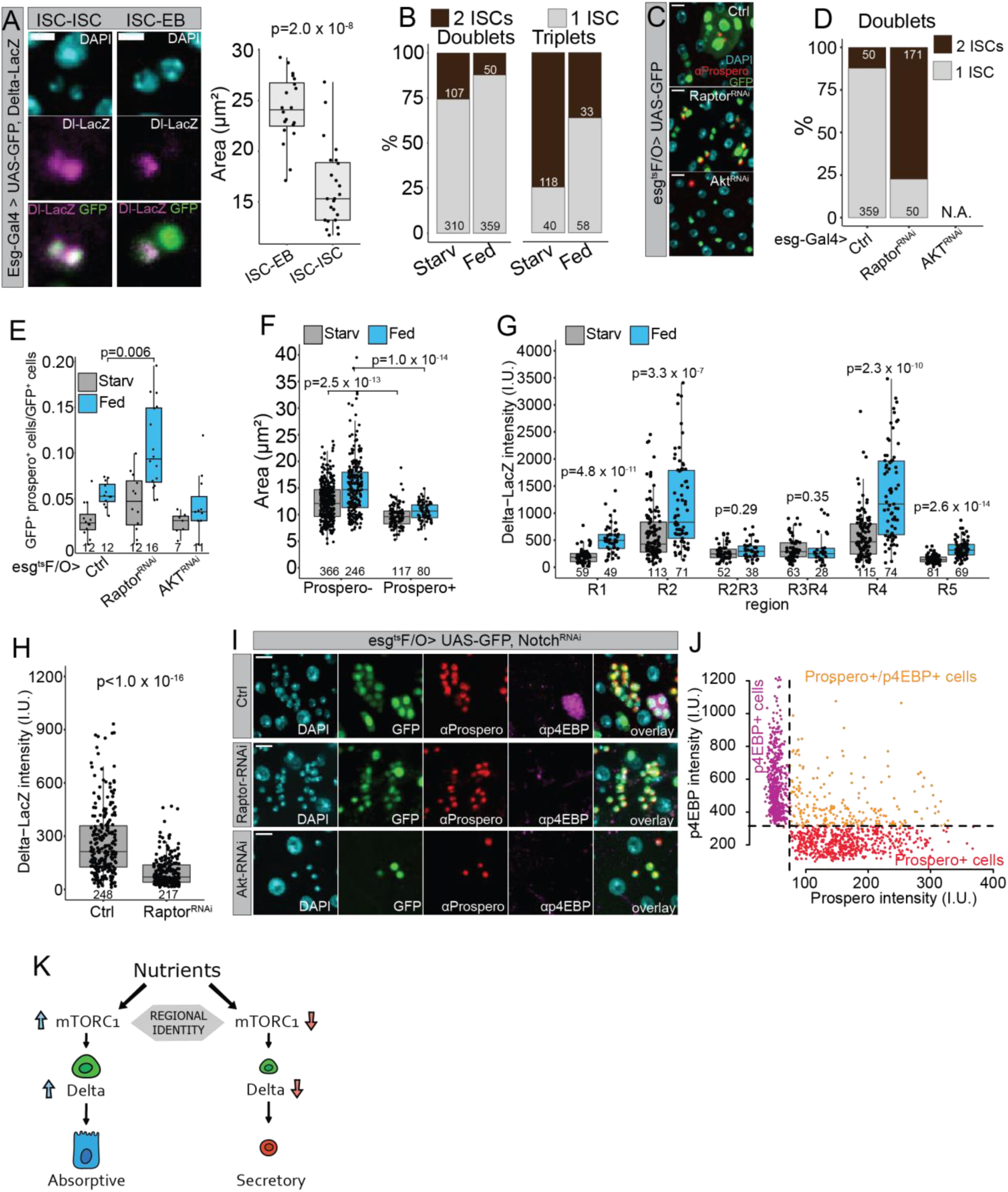
ISC fate is directed by cell autonomous mTORC1 activity. **A)** ISC size is associated with cell division mode. Representative images of an asymmetric ISC-EB doublet, and ISC-ISC doublet from midguts of female flies of genotype esg-Gal4>UAS-GFP, Delta-LacZ kept in holidic diet and stained by αβ-Galactosidase (magenta), and DAPI (cyan). Scale bar 10 μm. Right panel: quantification of ISC nucleus area from ISC-EB and ISC-ISC doublets. Pooled data from N=2 midguts from the R4b region. **B-E)** Feeding and mTORC1 activity regulates ISC fate. **B)** Delta-LacZ positive ISCs in esg^+^ cell doublets and triplets from midguts of female flies kept in either starvation or holidic diet. Pooled data from N^starved^=3 and N^fed^=4 midguts from the R4b region. N^doublets^ and N^triplets^ are indicated in the figure panel. Detailed experimental design of the experiment is depicted in Figure S2D. **C)** Representative images from midguts of female flies of genotype esg^ts^F/O>UAS-GFP (Ctrl) in combination with Raptor-RNAi or Akt-RNAi. Flies were kept in holidic diet for seven days and midguts were stained by αProspero (red) and DAPI (cyan). Scale bar 10 μm. Experimental design as in Figure 1A. **D)** Delta-LacZ positive ISCs in esg^+^ cell doublets from midguts of female flies of genotype esg-Gal4>fly-FUCCI, Delta-LacZ (Ctrl) in combination with Raptor-RNAi or Akt-RNAi. Pooled data from N^Ctrl^=4 and N^raptor-RNAi^=5 midguts from the R4b region. N^doublets^ are indicated in the figure panel. Detailed experimental design of the experiment is depicted in Figure S2D. **E)** Relative number of GFP positive and αProspero positive cells from the experiment depicted in C. Quantifications are performed from the R4b region. N^guts^ are indicated in the figure panel. Pooled data from two independent experiments. **F)** Prospero^+^ ISCs are smaller compared to Prospero-ISCs. Nuclear area of αProspero- and αProspero^+^ cells from starved and fed midgut. Quantifications are performed, and pooled, from R1, R2, borders flanking R3, R4 and R5. **G)** Delta expression is region-specific. Delta-LacZ intensity measurements (αβ-Galactosidase immunostaining) from R1, R2, borders flanking R3, R4 and R5 from midguts of female flies kept in either starvation or holidic diet. Experimental design as in Figure 1A. **H)** Delta expression is promoted by physiological mTORC1 activity. Delta-LacZ intensity measurements (αβ-Galactosidase immunostaining) from midguts of female flies of genotype esg-Gal4>fly-FUCCI, Delta-LacZ (Ctrl) in combination with Raptor-RNAi. Quantifications are performed from the R4b region. Pooled data from N^Ctrl^=4 and N^raptor-RNAi^=4 kept in holidic diet. N^cells^ are indicated in the figure panel. Experimental design as in Figure S2D. **I & J)** Physiological mTORC1 activity directly inhibits EE cell fate. **I)** Representative images from midguts of female flies of genotype esg^ts^F/O>UAS-GFP, Notch-RNAi in combination with Raptor-RNAi or Akt-RNAi. Flies were kept in holidic diet and midguts were stained by αProspero (red), αp4EBP (magenta) and DAPI (cyan). Scale bar 10 μm. Experimental design as in Figure 1A. **J)** αProspero and αp4EBP intensity dependency from GFP positive nuclei of female flies of genotype esg^ts^F/O>UAS-GFP, Notch-RNAi. Quantifications are performed from the R4c-R5 regions. Experimental design as in Figure 1A. **K)** A model depicting the role of nutrients and mTORC1 signaling in directing ISC fate during intestinal adaptation. p values in **A, E, F, G** and **H** were obtained by Wilcoxon rank-sum test with multiple testing correction (FDR<0.05) (in **G**).

To directly address the causal relationship between mTORC1-mediated ISC growth and cell fate we monitored esg^ts^F/O clones expressing RNAi against Raptor in starved and fed flies. Despite the observed growth inhibition upon ISC Raptor knockdown (Figure 2F), the ISC-derived clones contained cell numbers comparable to control clones (Figure 4C, S2B). This was in contrast to the knockdown of AKT, which strongly inhibited the feeding-induced increase in numbers of ISC-derived cells (Figure 4C, S2B), consistent with the important role of PI3K/AKT signaling in ISC proliferation^9,11^. As the knockdown of Raptor did not prevent ISC division, we were able to address the role of mTORC1 activity in symmetric vs. asymmetric division. Consistent with the indirect evidence associating increased ISC size with asymmetric ISC-EB division, inhibition of mTORC1 signaling significantly reduced the formation of ISC-EB doublets, increasing the relative amount of symmetric ISC-ISC doublets (Figure 4D, S2D). Thus, high mTORC1 activity primes the ISCs for asymmetric division to produce more absorptive cells.

In addition to EBs, ISCs can differentiate into small diploid EE cells. EE cells are marked by Prospero expression and they can arise through three mechanisms: by asymmetric ISC-EE division, symmetric ISC division producing two EE cells or by direct ISC to EE differentiation^28^. Notably, Prospero positive EEs do not arise through a Su(H)-positive EB intermediate in the adult posterior midgut^28^. The asymmetric ISC-EE doublet was very rare in our conditions (not shown). Interestingly, knockdown of Raptor by the esg^ts^F/O driver led to significantly increased number of EE cells in R4 region compared to the control clones under fed conditions (Figure 4C, E).

This result suggests that in addition to the symmetric ISC-ISC divisions, low mTORC1 signaling and small ISC size is associated with increased frequency of EE cell differentiation.

Since the EB vs. EE fate determination is controlled by the activity of Delta-Notch signaling^6,29,30^, we wanted to explore the relationship between ISC growth and Delta expression. ISCs with low Delta expression can differentiate into the EE fate through Prospero^+^ pre-EE state^28,31^. Consistently, we noticed that Delta^+^ Prospero^+^ pre-EE cells are expressing significantly less Delta as compared to the Delta^+^ Prospero-ISCs (Figure S2C). Interestingly, these pre-EE cells were also significantly smaller, compared to the rest of the Delta^+^ ISC population (Figure 4F). To further test the correlation between ISC size and Delta expression, we measured the regional Delta-LacZ intensity in starved and fed animals. Remarkably, Delta expression is highly region dependent, being low in the regions with small ISCs (R1, borders flanking R3 and R5), and high in the regions with large ISCs (R2 and R4) (Figure 4G). Furthermore, the expression of Delta is diet-dependent, being most strongly feeding activated in R2 and R4 and insensitive to feeding in both R3 borders (Figure 4G). Given the strong regional correlation with Delta expression, ISC size and differentiation, we asked if Delta expression is mTORC1 dependent. Indeed, Raptor knockdown in ISCs significantly reduced Delta-LacZ intensity in R4 (Figure 4H). Taken together, our results are consistent with a model that large ISCs with high mTORC1 activity express high levels of Delta, which directs ISC differentiation towards EB fate in a region-specific manner.

Elevated EE differentiation upon TORC1 inhibition (Figure 4E) might reflect direct inhibition of EE differentiation by mTORC1 or be an indirect consequence of reduced EB differentiation. To distinguish between these alternatives, we investigated the relationship between mTORC1 signaling and EE cell differentiation in the context of *Notch* loss-of-function (LOF), which inhibits ISC differentiation towards the EB lineage^29,30^. *Notch* LOF leads to the formation of proliferative endocrine progenitor tumors with a mixed population of small Pros^+^ and Pros-cells^32^. We stained *Notch* LOF tumors in fed conditions with the mTORC1 activity marker anti-p4EBP, which led to the identification of p4EBP^-^ and p4EBP^+^ subclusters (Figure 4I). Strikingly, Prospero expression was found only in p4EBP^-^, while the p4EBP^+^ cell clusters were devoid of Prospero expression (Figure 4J). The strong anti-correlation between mTORC1 activity and Prospero expression implied that high mTORC1 signaling directly inhibits EE cell fate. This was indeed the case, as Raptor knockdown in the *Notch* LOF clones abolished the Prospero^-^ p4EBP^+^ clusters, turning the tumour clusters homogenous for Prospero^+^ cells (Figure 4I). In contrast, knockdown of Akt completely prevented the tumorous growth of the *Notch* LOF clones (Figure 4I), consistent with its anti-proliferative effect (Figure 4C, S2B). In conclusion, our data shows that nutrient-regulated mTORC1 signaling directs ISCs towards the asymmetric ISC-EB division, while directly inhibiting EE cell differentiation (Figure 4K).

### Nutrition affects aging-induced deregulation of ISC differentiation

Aging is known to cause intestinal dysplasia manifested by the accumulation of mis-differentiated cells leading to progressive loss of epithelial homeostasis^33,34^. Considering the coupling of ISC fate determination and nutrient sensing, we asked if deregulated ISC differentiation in the aging intestine can be influenced by nutrition. To this end, we compared the intestines of aged *ad libitum* fed flies to flies of the same age exposed to repeated feeding-fasting cycles (intermittent fasting) (Figure S3A). As reported previously, we found that the midguts of aged *ad libitum* fed flies accumulate EEs and ISCs^34^ (Figure 5A). Strikingly, in the midguts of intermittent fasted flies the abnormal accumulation of EEs and ISCs was significantly suppressed (Figure 5A, S3B-D). These results suggests that in the aged *ad libitum* fed flies ISCs have progressively lost the ability to respond to nutrients, and hence favor symmetric division and EE cell differentiation over asymmetric ISC-EB divisions. To test this hypothesis, we exposed the old *ad libitum* fed flies to starvation and refeeding cycle and compared the intestinal adaptation response to the intermittent fasted flies of the same age. Interestingly, the feeding-induced increase in the cell number was very modest, indicating that the ISCs are not proliferating as a response to the fed state. In contrast, midguts of aged intermittent fasted flies better maintain their ability to increase intestinal cell numbers upon refeeding (Figure 5B, C). This is consistent with the more adaptive growth in the intermittent fasted flies. Hence, our results imply that aging induces a progressive decline in ISC nutrient activation while intermittent fasting can delay this process.

**Figure 5.**
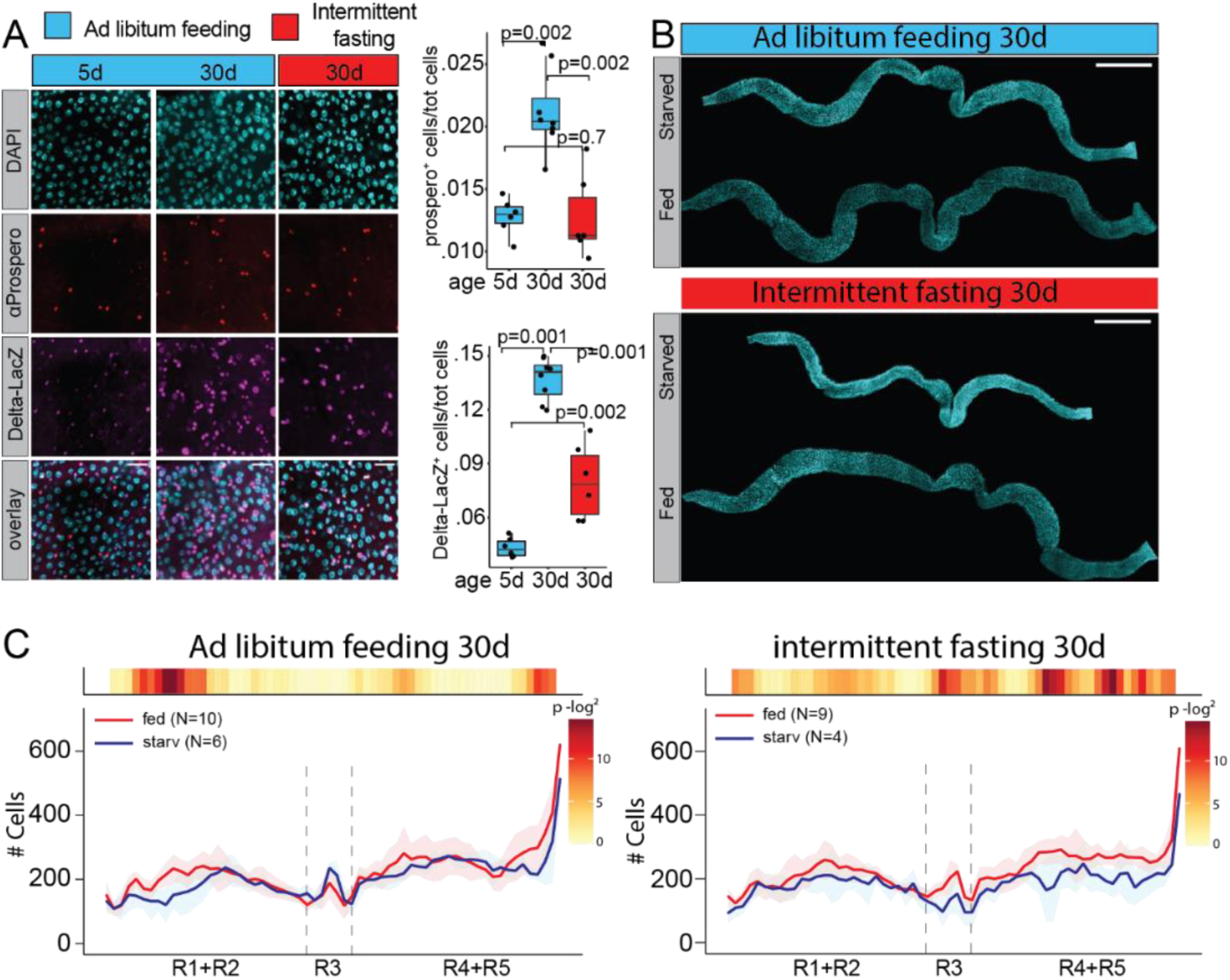
Lifelong intermittent fasting protects ISCs from aging induced decline of nutrient sensing. **A)** Lifelong intermittent fasting reduce the accumulation of EE cells in aging midgut. Representative images of αProspero stained midguts from the R4b region, and quantification of αProspero and Delta-LacZ positive cells from the same region. Scale bar 25 μm. Experimental design as in Figure S3A. **B)** Representative images of starved and refed midguts after 30 days of ad libitum feeding or intermittent fasting. Scale bar 500 μm. **C)** Regional quantification of starved versus refed midgut cell numbers along the the A/P axis after 30 days of *ad libitum* feeding or intermittent fasting. p values in **A** were obtained by Wilcoxon rank-sum test with multiple testing correction (FDR<0.05). p values in **A** were obtained by Wilcoxon rank-sum test with multiple testing correction (FDR<0.05). p values in **C** were obtained by Wilcoxon rank-sum test using continuity and multiple testing correction (FDR<0.05. See also Figure S3.

## DISCUSSION

Organ-wide analysis of the *Drosophila* midgut nutrient adaptation identified regionally distinct patterns of EC growth, ISC proliferation and fate determination upon transition from fasted to fed state. Fed animals display activation of the mTORC1 nutrient sensing pathway in ECs or ISCs in a region-specific manner. The ISC mTORC1 activity is coupled to cell cycle progression, being highest during S and G2/M phases. Consistent with the activation of mTORC1, ISC size is prominently increased already one day after the transition to a fed state. Large ISCs with high mTORC1 activity display high expression of the Notch ligand Delta and are directed towards asymmetric division to produce an ISC-EB pair, thus promoting differentiation towards the absorptive lineage. Furthermore, mTORC1 activity inhibits differentiation towards the secretory EE lineage, even under conditions where differentiation to the absorptive lineage is inhibited (Notch loss-of-function). Thus, our results show that dynamic control of ISC size allows coupling of nutrient-responsive mTORC1 signaling to cell fate determination, regulating regionalized ISC differentiation patterns during adaptive growth of the intestine.

Previous work have shown that feeding promotes *Drosophila* ISC proliferation and EC size as well as villi length and cellular turnover of the mouse small intestine^2,4,10,24,35,36^. Our quantitative organ-wide analysis of the *Drosophila* midgut revealed substantial regional differences in EC growth and ISC proliferation. Moreover, our data uncovered dynamic changes in regional distributions of ISC differentiation. What are the physiological reasons for the regional differences in the adaptive responses? Enterocytes in different regions differ in their metabolic functions^12,14,37,38^. Such differences might lead to locally different EC turnover rate, determining the need for ISC proliferation/differentiation upon intestinal nutrient adaptation. In addition to ECs, EE cells can be divided into regionally distributed subtypes depending on the hormones they express^38,39^. It will be interesting to explore whether the observed increase in EE cell numbers in the middle parts of the midgut is associated with feeding-induced changes in specific EE cell subtypes. Mechanistically, we observed that transition to a fed state activated the mTORC1 signaling pathway throughout the whole intestine, but with distinct local distributions between the ISCs vs. ECs. How are the regional patterns of ISC mTORC1 activity achieved? As ISCs are located on the basal side of epithelium, their access to intestinal lumen-derived nutrients likely depends on their local interactions with the nutrient-absorbing ECs. In line with this idea is the finding that the metabolism of mouse ISCs are regulated by controlled exchange of nutrients with their neighboring cells^40^. We observed mutually exclusive regional activation patterns of the mTORC1 between ISCs and ECs which is consistent with a model that ECs might either activate a cell autonomous growth program, or alternatively, channel the mTORC1 activating nutrients to the neighboring stem cells, facilitating their proliferation and growth. Furthermore, it has been shown that the EC-ISC contact restrict ISC divisions through E-cadherin mediated signaling, and that enlarged ECs, through expression of a constitutively active Insulin receptor, inhibit the proliferation of nearby ISCs^10,41^. Moreover, an inverse correlation between ISC proliferative activity and the EC cell size was observed by screening infection response of genetically different *Drosophila* lines^42^. Future studies should be directed to resolve the role of local tissue environment in coordinating ISC proliferation, growth and differentiation as well as the physiological roles of region-specific ISC regulation.

Previous work has demonstrated that mTORC1 contributes to activation of quiescent somatic stem cells^21^, while prolonged superphysiological mTORC1 activity in stem cells result in loss of stemness and self-renewal^18,19^. Our quantitative analysis revealed that dynamic regulation of ISC size through mTORC1 signaling serves as an early fate determinant, directing the outcome of stem cell activation upon intestinal nutrient adaptation. How does ISC size define the outcome of fate decision? Notch signaling pathway activity is the key determinant between the absorptive vs. secretory lineages. The level of Delta expression dictates the future fate of dividing ISC, as high Delta-induced Notch activation after asymmetric division promotes the absorptive lineage differentiation^6^. We found that the expression of Delta is strongly regulated by dietary nutrients in an mTORC1 dependent manner, specifically in regions displaying ISC growth and consequently increased number of EBs. While it remains unresolved, how ISC size influences Delta expression, it is tempting to speculate that cell size impacts the perception of niche derived signals, and consequently, the fate of the dividing ISC. *Drosophila* ISCs reside basally in a space restricted by the neighboring ECs, the basement membrane and the underlying muscle layer^5^. The ISC contact with the basement membrane through integrin signaling is essential for the maintenance of stem cell identity^6,43^. During nutrient induced midgut growth, symmetric ISC divisions take place in a basal-to-basal orientation whereas the asymmetric ISC-EB divisions in a basal-to-apical orientation^2^. ISC growth-induced changes in the relative contact surface to the basement membrane could potentially alter the niche derived signaling strength, and hence, the outcome of the daughter cell fate. In line with this idea, it was recently shown that ISC shape, which is linked to its surface to volume ratio, changed the strength of the niche derived intracellular signaling activities of the ISC in mouse small intestine^44^.

The role of nutrient sensing in the regulation of somatic stem cell function, and consequently tissue homeostasis, is of central importance in physiological adaptation to changing nutrient conditions. Our results on the regulation of ISC size explain how nutrient induced signaling can control locally ISC differentiation, to achieve region-specific adaptive responses. The coupling between nutrient sensing and cell fate may open new intervention possibilities to delay aging-induced tissue decline, manifested as deregulated ISC differentiation. Our findings on the beneficial effects of intermittent fasting on ISC differentiation, together with the role of stem cell size as an aging factor^20,45^ and the impact of ISC mTORC1 signaling in aging^21^, highlight the need for more detailed investigation on the role of nutrient-induced control of ISC size in the context of aging tissue.

## MATERIALS & METHODS

### Drosophila stocks and husbandry

Fly stocks used in this study: w; esg-Gal4, Tub-Gal80-ts, UAS-GFP ; UAS-Flp, Act>CD2>Gal4 (esg^ts^F/O)^25^, w; esg-Gal4, Tub-Gal80^ts^, UAS-GFP (esg^ts^)^46^, Delta-Gal4^47^, Delta-LacZ (Dl-LacZ, Bloomington 11651), Gbe+Su(H)-lacZ (Su(H)-LacZ,^48^), Raptor-RNAi (Bloomington 31529), Akt-RNAi (Bloomington 33615), Notch-RNAi (Bloomington 27988), Fly-Fucci^26^, UAS-Reaper (Bloomington 5824). Flies were maintained at 25°C, on medium containing agar 0.6% (w/v), malt 6.5% (w/v), semolina 3.2% (w/v), baker’s yeast 1.8% (w/v), nipagin 2.4%, propionic acid 0.7%.

### Immunohistochemistry

For immunofluorescence staining, intestines were dissected in PBS and fixed in 8% paraformaldehyde for 3 hours. Tissues were washed with 0.1% Triton-X 100 in PBS and blocked in 1% bovine serum albumin for 1 h. Subsequently, tissues were stained with anti-β-Galactosidase (1:400) (MP Biomedicals cat no: 0855976-CF), anti-Prospero (1:1000) (MR1A, DSHB), anti-p4EBP (CST 2855) antibodies. The samples were mounted in Vectashield mounting media with DAPI (Vector Laboratories) and imaged using the Aurox clarity confocal system (Aurox). Images were further processed by the ImageJ software.

### FACS and RNA-Seq

Gut dissection and FACS dissociation was essentially performed as in Dutta et al., 2015 with the following modifications. 80-100 guts per sample were dissociated in 4 mg/ml Elastase with pipetting 15-20X every 15 minutes for 1 hour at 28ºC with shaking at 600 rpm. After dissociation, cells were filtered through a 40μM filter and FACS-sorted based on CFP/YFP and mRFP-signal. Three triplicate samples of 80-100 guts each were used to sort G1 and G2 cells into RNA isolation buffer. RNA was isolated using the Arcturus PicoPure RNA isolation kit (ThermoFischer) and subsequently amplified using the RiboAmp HS-Plus kit (ThermoFischer). Libraries were generated using the TruSeq Stranded mRNA Library Prep Kit (Illumina) and subsequently sequenced as 50 bp single-end on an Illumina HiSeq2500. Reads were mapped using Bowtie and reads were counted using htseqcount. Tables of raw counts per gene/sample were analyzed with the R package DESeq2 for differential expression^49^. G1 and G2 populations were compared with each other. Genes with an adjusted p-value <0.05 were considered to be differentially expressed (DE). Volcano plots were generated by VolcanoseR^50^. The raw sequencing data is deposited into the Gene Expression Omnibus (GEO) under the accession number GSE222254.

### Microscopy and image processing

Fixed and immunostained midguts were mounted in between a microscope slide with 0.12μm spacers and a coverslip, followed by imaging by the Aurox clarity spinning disc confocal microscope. Images were segmented by Stardist and raw feature data were exported and used as input for LAM for further analysis^16^.

### Statistical analysis

Statistical analyses were performed in R/Bioconductor. For parametric data, two-sample t-test or two-way ANOVA in conjunction with Tukey’s HSD test was used. For the non-parametric data Wilcoxon rank-sum test with multiple testing correction (FDR<0.05) was used. p-values for the RNAseq experiment were calculated using the Wald significance test and adjusted with Benjamini & Hochberg correction for multiple testing.

## Supporting information

Supplementary images

## ACKNOWLEDGEMENTS

We thank Jette Lengefeld and Nalle Pentinmikko for providing feedback on the manuscript. We thank Bloomington *Drosophila* Stock Center, Norman Zielke, Minna Poukkula and Bruce Edgar for providing fly stocks, and Kathrin Schubert, Maria Locke and Marco Groth from the Flow Cytometry and Next Generation Sequencing Facilities at the FLI-Leibniz Institute on Aging for expert technical assistance. Funding was provided by the Academy of Finland through the MetaStem Center of Excellence funding to V.H. (312439) and a project grant (137530) to J.M., Sigrid Juselius Foundation (V.H., J.M.), Erkko Foundation (V.H.), Novo Nordisk Foundation (NNF19OC0057478 to V.H.), and DFG research grant number (KO5594/1-1 to J.K.). This study was facilitated by the University of Helsinki *Drosophila* core facility (Hi-Fly) and the Light microscopy unit (LMU) supported by Biocenter Finland and Helsinki Institute of Life Science.

## AUTHOR CONTRIBUTIONS

Conceptualization, J.M. and V.H.; Methodology, V.H., J.M., J.K., and A.V.; Investigation, J.M., A.V., G.F., J.K.; Data Curation, J.M., A.V., G.F., J.K.; Writing – Original Draft, J.M., and V.H.;

Writing – Review & Editing, J.M., A.V., G.F., J.K., and V.H.; Visualization, J.M., G.F., J.K., and

A.V.; Supervision, J.M., V.H.; Funding Acquisition, V.H., J.K., and J.M.

## DECLARATION OF INTERESTS

The authors declare no competing interests.

## Notes

### Competing Interest Statement

The authors have declared no competing interest.

## REFERENCES

1. Goodlad, R. A. & Wright, N. A. The effects of starvation and refeeding on intestinal cell proliferation in the mouse. Virchows Archiv B Cell Pathol 45, 63–73 (1984).

2. O’Brien, L. E., Soliman, S. S., Li, X. & Bilder, D. Altered Modes of Stem Cell Division Drive Adaptive Intestinal Growth. Cell 147, 603–614 (2011).

3. Stojanović, O., Miguel-Aliaga, I. & Trajkovski, M. Intestinal plasticity and metabolism as regulators of organismal energy homeostasis. Nat Metab (2022) doi:10.1038/s42255-022-00679-6.

4. Bonfini, A. et al. Multiscale analysis reveals that diet-dependent midgut plasticity emerges from alterations in both stem cell niche coupling and enterocyte size. e Life 10, e64125 (2021).

5. Miguel-Aliaga, I., Jasper, H. & Lemaitre, B. Anatomy and Physiology of the Digestive Tract of Drosophila melanogaster. Genetics 210, 357–396 (2018).

6. Ohlstein, B. & Spradling, A. Multipotent Drosophila Intestinal Stem Cells Specify Daughter Cell Fates by Differential Notch Signaling. Science 315, 988–992 (2007).

7. Perdigoto, C. N., Schweisguth, F. & Bardin, A. J. Distinct levels of Notch activity for commitment and terminal differentiation of stem cells in the adult fly intestine. Development 138, 4585–4595 (2011).

8. Obniski, R., Sieber, M. & Spradling, A. C. Dietary Lipids Modulate Notch Signaling and Influence Adult Intestinal Development and Metabolism in Drosophila. Developmental Cell (2018) doi:10.1016/j.devcel.2018.08.013.

9. Kapuria, S., Karpac, J., Biteau, B., Hwangbo, D. & Jasper, H. Notch-Mediated Suppression of TSC2 Expression Regulates Cell Differentiation in the Drosophila Intestinal Stem Cell Lineage. PLoS Genetics 8, e1003045 (2012).

10. Choi, N. H., Lucchetta, E. & Ohlstein, B. Nonautonomous regulation of Drosophila midgut stem cell proliferation by the insulin-signaling pathway. Proceedings of the National Academy of Sciences 108, 18702–18707 (2011).

11. Mattila, J., Kokki, K., Hietakangas, V. & Boutros, M. Stem Cell Intrinsic Hexosamine Metabolism Regulates Intestinal Adaptation to Nutrient Content. Developmental Cell (2018) doi:10.1016/j.devcel.2018.08.011.

12. Buchon, N. et al. Morphological and Molecular Characterization of Adult Midgut Compartmentalization in Drosophila. Cell Reports 3, 1725–1738 (2013).

13. Gebert, N. et al. Region-Specific Proteome Changes of the Intestinal Epithelium during Aging and Dietary Restriction. Cell Reports 31, 107565 (2020).

14. Marianes, A. & Spradling, A. C. Physiological and stem cell compartmentalization within the Drosophila midgut. eLife 2, e00886 (2013).

15. Dutta, D. et al. Regional Cell-Specific Transcriptome Mapping Reveals Regulatory Complexity in the Adult Drosophila Midgut. Cell Rep 12, 346–358 (2015).

16. Viitanen, A. et al. An image analysis method for regionally defined cellular phenotyping of the Drosophila midgut. Cell Reports Methods 1, 100059 (2021).

17. Saxton, R. A. & Sabatini, D. M. mTOR Signaling in Growth, Metabolism, and Disease. Cell 168, 960–976 (2017).

18. Castilho, R. M., Squarize, C. H., Chodosh, L. A., Williams, B. O. & Gutkind, J. S. mTOR Mediates Wnt-Induced Epidermal Stem Cell Exhaustion and Aging. Cell Stem Cell 5, 279–289 (2009).

19. Amcheslavsky, A., Ito, N., Jiang, J. & Ip, Y. T. Tuberous sclerosis complex and Myc coordinate the growth and division of Drosophila intestinal stem cells. J. Cell Biol. 193, 695–710 (2011).

20. Lengefeld, J. et al. Cell size is a determinant of stem cell potential during aging. Sci. Adv. 7, eabk0271 (2021).

21. Haller, S. et al. mTORC1 Activation during Repeated Regeneration Impairs Somatic Stem Cell Maintenance. Cell Stem Cell 21, 806-818.e5 (2017).

22. Rodgers, J. T. et al. mTORC1 controls the adaptive transition of quiescent stem cells from G0 to GAlert. Nature 510, 393–396 (2014).

23. Yilmaz, Ö. H. et al. mTORC1 in the Paneth cell niche couples intestinal stem-cell function to calorie intake. Nature 486, 490–495 (2012).

24. McLeod, C. J., Wang, L., Wong, C. & Jones, D. L. Stem Cell Dynamics in Response to Nutrient Availability. Current Biology 20, 2100–2105 (2010).

25. Jiang, H. et al. Cytokine/Jak/Stat signaling mediates regeneration and homeostasis in the Drosophila midgut. Cell 137, 1343–1355 (2009).

26. Zielke, N. et al. Fly-FUCCI: A versatile tool for studying cell proliferation in complex tissues. Cell Rep 7, 588–598 (2014).

27. Pallares-Cartes, C., Cakan-Akdogan, G. & Teleman, A. A. Tissue-Specific Coupling between Insulin/IGF and TORC1 Signaling via PRAS40 in Drosophila. Developmental Cell 22, 172–182 (2012).

28. Zeng, X. & Hou, S. X. Enteroendocrine cells are generated from stem cells through a distinct progenitor in the adult Drosophila posterior midgut. Development 142, 644–653 (2015).

29. Micchelli, C. A. & Perrimon, N. Evidence that stem cells reside in the adult Drosophila midgut epithelium. Nature 439, 475–479 (2006).

30. Ohlstein, B. & Spradling, A. The adult Drosophila posterior midgut is maintained by pluripotent stem cells. Nature 439, 470–474 (2006).

31. Chen, J. et al. Transient Scute activation via a self-stimulatory loop directs enteroendocrine cell pair specification from self-renewing intestinal stem cells. Nat Cell Biol 20, 152–161 (2018).

32. Patel, P. H., Dutta, D. & Edgar, B. A. Niche appropriation by Drosophila intestinal stem cell tumours. Nat. Cell Biol. 17, 1182–1192 (2015).

33. Biteau, B., Hochmuth, C. E. & Jasper, H. JNK Activity in Somatic Stem Cells Causes Loss of Tissue Homeostasis in the Aging Drosophila Gut. Cell Stem Cell 3, 442–455 (2008).

34. Choi, N.-H., Kim, J.-G., Yang, D.-J., Kim, Y.-S. & Yoo, M.-A. Age-related changes in Drosophila midgut are associated with PVF2, a PDGF/VEGF-like growth factor. Aging Cell 7, 318–334 (2008).

35. Altmann, G. G. Influence of starvation and refeeding on mucosal size and epithelial renewal in the rat small intestine. American Journal of Anatomy 133, 391–400 (1972).

36. Dunel-Erb, S. et al. Restoration of the jejunal mucosa in rats refed after prolonged fasting. Comp. Biochem. Physiol., Part A Mol. Integr. Physiol. 129, 933–947 (2001).

37. Aliluev, A. et al. Diet-induced alteration of intestinal stem cell function underlies obesity and prediabetes in mice. Nat Metab 3, 1202–1216 (2021).

38. Haber, A. L. et al. A single-cell survey of the small intestinal epithelium. Nature 551, 333–339 (2017).

39. Chen, J., Kim, S.-M. & Kwon, J. Y. A Systematic Analysis of Drosophila Regulatory Peptide Expression in Enteroendocrine Cells. Mol Cells 39, 358–366 (2016).

40. Rodríguez-Colman, M. J. et al. Interplay between metabolic identities in the intestinal crypt supports stem cell function. Nature 543, 424–427 (2017).

41. Liang, J., Balachandra, S., Ngo, S. & O’Brien, L. E. Feedback regulation of steady-state epithelial turnover and organ size. (2017) doi:10.1101/161943.

42. Tamamouna, V. et al. Evidence of two types of balance between stem cell mitosis and enterocyte nucleus growth in the Drosophila midgut. Development 147, dev189472 (2020).

43. Lin, G. et al. Integrin signaling is required for maintenance and proliferation of intestinal stem cells in Drosophila. Dev. Biol. 377, 177–187 (2013).

44. Pentinmikko, N. et al. Cellular shape reinforces niche to stem cell signaling in the small intestine. Sci Adv 8, eabm1847 (2022).

45. Davies, D. M., van den Handel, K., Bharadwaj, S. & Lengefeld, J. Cellular enlargement - A new hallmark of aging? Front. Cell Dev. Biol. 10, 1036602 (2022).

46. Jiang, H. & Edgar, B. A. EGFR signaling regulates the proliferation of Drosophila adult midgut progenitors. Development 136, 483–493 (2009).

47. Zeng, X., Chauhan, C. & Hou, S. X. Characterization of midgut stem cell- and enteroblast-specific Gal4 lines in drosophila. genesis 48, 607–611 (2010).

48. Furriols, M. & Bray, S. A model Notch response element detects Suppressor of Hairless–dependent molecular switch. Current Biology 11, 60–64 (2001).

49. Love, M. I., Huber, W. & Anders, S. Moderated estimation of fold change and dispersion for RNA-seq data with DESeq2. Genome Biol 15, 550 (2014).

50. Goedhart, J. & Luijsterburg, M. S. VolcaNoseR is a web app for creating, exploring, labeling and sharing volcano plots. Sci Rep 10, 20560 (2020).

