## Supplementary images for "Stem cell growth directs region-specific cell fate decisions during intestinal nutrient adaptation"

Jaakko Mattila et al.

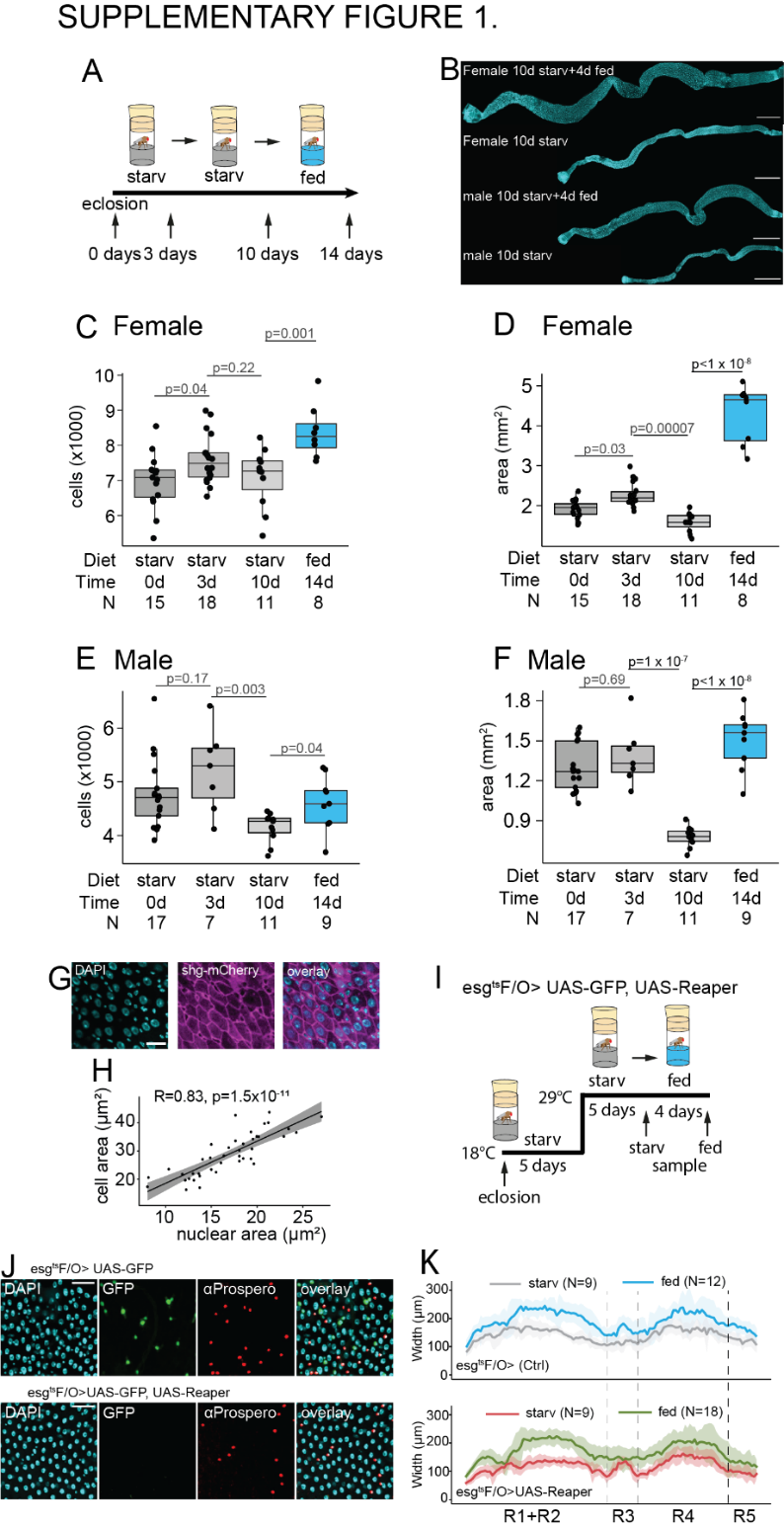

**Supplementary Figure 1.** Related to the main Figure 1.

**A-F)** *Drosophila* midgut cell number and size is dynamically regulated by feeding.  **A)** Experimental design used to obtain data in panels **B-F**. Age matched, mated wild type (OregonR) females and males were kept at +25°C in starvation for ten days, and then shifted to the holidic diet for an additional 4 days. Midgut samples were obtained at 0, 3, 10 and 14 days. **B)** Representative images of female and male midguts at 10 (starved) and 14 days (fed 4 days) time points. Scale bar in is 200µm **C)** Quantification of female midgut cell numbers at the indicated time points. Quantified from one cell layer of a flattened, two-layer midgut. Hence the total midgut cell number is approximately 2x of the indicated value. **D)** Quantification of total surface area of a flattened female midgut at the indicated time points. **E)** Quantification of male midgut cell numbers at the indicated time points. Quantified from one cell layer of a flattened, two-layer midgut. Hence the total cell number is approximately 2x of the indicated value. **F)** Quantification of total surface area of a flattened male midgut at the indicated time points. **G & H)** Midgut nuclear area and cell area correlate. **G)** Representative images of shg-mCherry (magenta) and DAPI (cyan) stained midguts. Scale bar in is 20µm. **H)** Spearman correlation between nucleus (DAPI stained nuclei in G) and cell (shg-mCherry restricted area in G) area. **I-K)** Dietary nutrient induced midgut growth is not dependent on ISCs. **I)** Experimental design used to obtain data in panel **J**. **J)** Representative images from midguts of female flies of genotypes esg^ts^F/O>UAS-GFP (Ctrl) or esg^ts^F/O>UAS-GFP, UAS-Reaper. **K)** Width profile along the A/P axis of starved and fed midguts of Ctrl (upper panel) and ISC depleted (lower panel) flies.

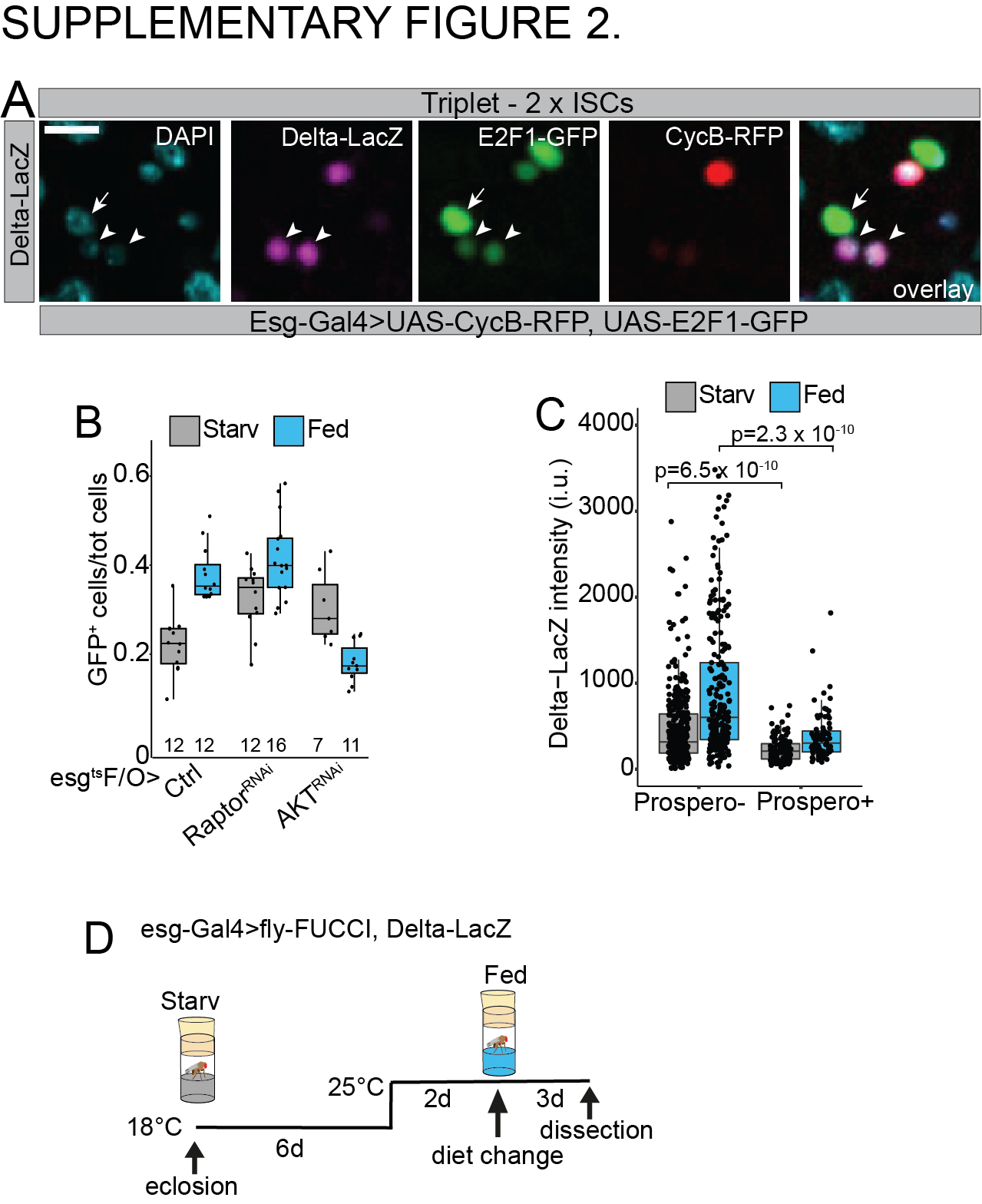

**Supplementary Figure 2.** Related to the main Figure 4.

**A)** Representative image of a cell triplet with two ISCs from female midgut of the genotype esg-Gal4>UAS-CycB-RFP, UAS-E2F1-GFP, Delta-LacZ. Arrowheads point to Delta-LacZ (magenta) positive ISCs and the arrow to esg-Gal4 expressing enteroblast. Scale bar 10µm. **B)** Quantification of relative GFP positive cell number from the experiment depicted in the main Figure 4C. Quantifications were performed from the R4bc region from midguts of female flies of genotype esg^ts^F/O>UAS-GFP (Ctrl) in combination with Raptor-RNAi or Akt-RNAi. **C)** Quantification of Delta-LacZ intensity (αβ-Galactosidase immunostaining) from αProspero positive EE cells from the experiment depicted in Figure 4G. Measurements were from R1, R2, borders flanking R3, R4 and R5 from midguts of female flies kept in either starvation or holidic diet. Experimental design as in Figure 1A. **D)** Experimental design used to obtain data in panels Figure 4B & 4D. Age matched, mated females of genotype esg-Gal4>UAS-CycB-RFP, UAS-E2F1-GFP, Delta-LacZ in combination with Raptor-RNAi or Akt-RNAi, were aged for six days, and then shifted to the holidic diet for 3 days. p values in C were obtained by Wilcoxon rank-sum test with multiple testing correction (FDR<0.05).

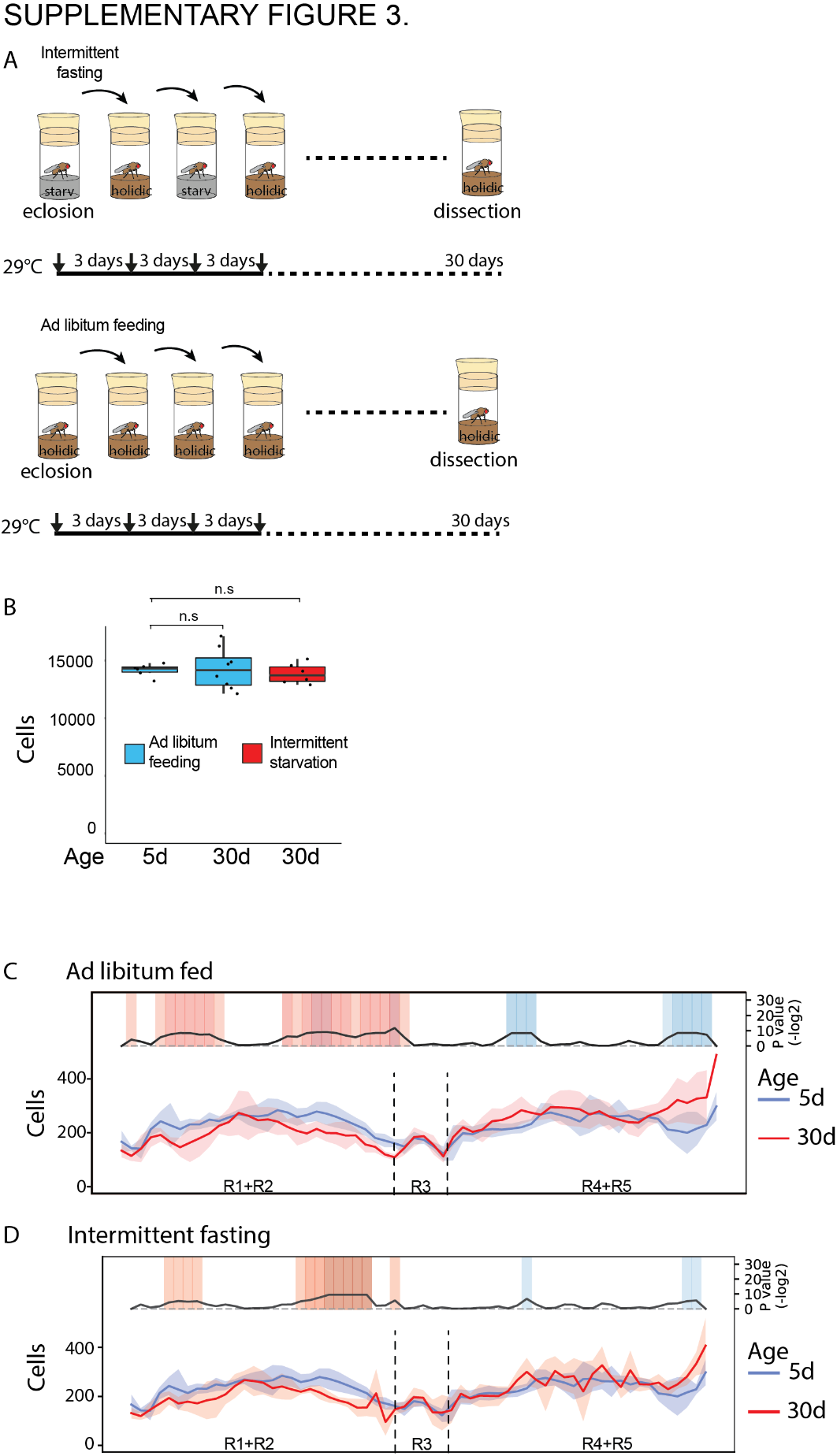

**Supplementary Figure 3.** Related to the main Figure 5.

**A)** Experimental design used to obtain data in Figure 5A. Age matched females of genotype esg-Gal4-ts>UAS-GFP, Delta-LacZ were kept at 29°C for 30 days in holidic diet (ad libitum fed) or flipped to fasting after three days feeding period (intermittent fasting). **B)** Quantification of total cell numbers from the experimented depicted in **A**. **C, D)** Total cell count comparisons along the midgut A/P axis between young (5d) and old (30d) **(C)** and young (5d) and old (30d) **(D)** intermittent fasted female midguts.
